# Identification and differentiation of *Pseudomonas* species in field samples using an *rpoD* amplicon sequencing methodology

**DOI:** 10.1101/2021.06.08.447643

**Authors:** Jonas Greve Lauritsen, Morten Lindqvist Hansen, Pernille Kjersgaard Bech, Lars Jelsbak, Lone Gram, Mikael Lenz Strube

## Abstract

Species of the genus *Pseudomonas* are used for several biotechnological purposes, including plant biocontrol and bioremediation. To exploit the *Pseudomonas* genus in environmental, agricultural or industrial settings, the organisms must be profiled at species level as their bioactivity potential differs markedly between species. Standard 16S rRNA gene amplicon profiling does not allow for accurate species differentiation. Thus, the purpose of this study was to develop an amplicon-based high-resolution method targeting a 760 nt region of the *rpoD* gene enabling taxonomic differentiation of *Pseudomonas* species in soil samples. The method was benchmarked on a sixteen membered *Pseudomonas* species mock community. All 16 species were correctly and semi-quantitatively identified using *rpoD* gene amplicons, whereas 16S rRNA V3V4 amplicon sequencing only correctly identified one species. We analysed the *Pseudomonas* profile in thirteen soil samples in northern Zealand, Denmark, where samples were collected from grassland (3 samples) and agriculture soil (10 samples). *Pseudomonas* species represented up to 0.7% of the microbial community, of which each sampling site contained a unique *Pseudomonas* composition. Thirty culturable *Pseudomonas* strains were isolated from each grassland site and ten from each agriculture site and identified by Sanger sequencing of the *rpoD* gene. In all cases, the *rpoD*-amplicon approach identified more species than found by cultivation, including hard-to-culture non-fluorescent pseudomonads, as well as more than found by 16S rRNA V3V4 amplicon sequencing. Thus, *rpoD* profiling can be used for species profiling of *Pseudomonas*, and large scale prospecting of bioactive *Pseudomonas* may be guided by initial screening using this method.

**Importance:** A high throughput sequence-based method for profiling of *Pseudomonas* species in soil microbiomes was developed and identified more species than 16S rRNA gene sequencing or cultivation. *Pseudomonas* species are used as biocontrol organisms and plant-growth promoting agents, and the method will allow tracing of specific species of *Pseudomonas* as well as enable screening of environmental samples for further isolation and exploitation.

## INTRODUCTION

*Pseudomonas* species are ubiquitous and can be isolated from a range of environments including plant rhizospheres, marine habitats, and animal tissues (1–4). Whilst the genus contains species that are pathogenic to plants and animals, several species express traits that enable their use in bioremediation, plant growth promotion or plant disease suppression (5–8). The underlying beneficial mechanisms are often linked to specific species or even strains, including the production of pathogen suppressing secondary metabolites, such as the antibiotic 2,4-diacetylphloroglucinol (DAPG) and pyoverdine siderophores (5, 9–11). Also, some strains promote growth of plants by solubilising inorganic nutrients such as phosphate and iron or by producing plant hormones (11–14). Members of the *Pseudomonas fluorescence* group, in particular, are a major source of bioactivity since they have markedly larger genomes than other pseudomonads (15) and a high number of biosynthetic gene clusters as determined by genomic analysis (16). In addition to strains expressing these and other beneficial traits, it is also becoming clear that the structure and diversity of the *Pseudomonas* community in bulk and rhizosphere soils *per se* can be associated with suppression of crop fungal pathogens (17, 18). Studies on the distribution, abundance and diversity of *Pseudomonas* spp. in soil and rhizosphere often rely on cultivation dependent analyses. However, Aagot *et al*. and others have demonstrated that cultivation of individual species of *Pseudomonas* is dependent on the specific conditions used (e.g. level of nutrients), and the decision of a specific cultivation medium is thus a source of bias (19). Given these biases, linking specific *Pseudomonas* species and/or community structures to certain ecosystem performance metrics (including suppression of crop fungal pathogens) remains a challenge.

Amplicon sequencing of the 16S rRNA gene has become the standard for culture-independent, taxonomic profiling of environmental microbial communities. However, the 16S rRNA gene is very similar across closely related *Pseudomonas* species with less than 1% nucleotide dissimilarity between many of the species (20). For example, in the subgroup of *P. putida*, the dissimilarities are between 0.16% and 2.31% (20). Therefore, 16S rRNA gene profiling only provides taxonomic resolution at genus level, and studies of *Pseudomonas* community structures and dynamics at species level require sequencing and analyses of other house-keeping genes. The *rpoD* gene, which encode the sigma 70 factor of RNA polymerase, is an excellent target gene for phylogenetic and taxonomic analyses of *Pseudomonas* species (21). Using a highly selective pair of *Pseudomonas rpoD* primers, PsEG30F-PsEG790R (21), an *rpoD*-amplicon sequencing method was used to analyse environmental DNA obtained from a single water sample (22). The method was developed for the 454/Roche GS-FLX platform and used an in-house *rpoD* database for sequence analysis. Given the discontinuation of the 454/Roche GS-FLX platform and the understanding of *Pseudomonas* phylogeny, there is a need for development of an amplicon-based method for reliable identification and differentiation of *Pseudomonas* species from environmental samples.

The purpose of the present study was to develop an amplicon sequencing protocol compatible with the Illumina MiSeq 300PE platform, and to establish a new and improved bioinformatic pipeline with an updated database build upon the *Pseudomonas* type strain collection and taxonomic framework from Hesse *et al*. (15). The *rpoD* amplicon method allowed *Pseudomonas* species differentiation in environmental soil samples and can guide future bioprospecting endeavours.

## RESULTS

### *In silico* target gene evaluation

We evaluated nine genes and their accompanying primer sets (14 in total) for their phylogenetic discriminatory power using *in silico* PCR against two sets of genomes: First, a library of the 166 type-strain genomes of Hesse *et al*. (15) acting as a well curated collection of all known *Pseudomonas* and their phylogeny, although with most genomes being in contig form. Second, a library of 465 genomes of *Pseudomonas* species available from NCBI, all of which are complete but with high redundancy and incomplete taxonomy (Table 1). The *rpoD* primer pair PsEG30F and PsEG790R (21) resulted in the best phylogenetic resolution along with the highest total number of individual *Pseudomonas* genomes amplified and the lowest total non-*Pseudomonas* amplifications. This pair amplified 160/166 (96.39%) of type strains and 460/465 (98.92%) of the complete genomes (Table S3) with no amplification of the negative controls. The *gyrB*-gene primers UP-1E / APrU showed 100% amplification of the type strains, but unfortunately also amplified 25% of the negative controls and had multiple instances of amplicons much longer than expected length. To further evaluate the phylogenetic resolution of the primers, a multiple sequence alignment of the amplicons and the resulting phylogeny was compared to the study by Hesse *et al*. (15), in which the 166 distinct *Pseudomonas* type strains were phylogenetically resolved based on Multiple Locus Sequencing Typing of 100 genes. Here, the *rpoD* primers PsEG30F and PsEG790R from Mulet *et al*. (21) produced amplicons with the closest similarity to the phylogenetic map of Hesse et al. (15), and separated all species in the phylogenetic tree. The primers generated a ~760 nt long amplicon of the *rpoD* gene, which unfortunately led to a sequencing gap of 160 nt using the 300PE platform. Therefore, two new reverse primers were designed, PsJL490R and PsJL628R, however, both had lower amplification of the type strain genomes at 89.16% and 83.73%, respectively. Moreover, the PsJL490R had poor *in vitro* amplification for the species of the synthetic community while the PsJL628R showed unspecific amplification of the negative controls *S. maltophila, A. xylosoxidans* and *A. brasilense* (Figure S1). Phylogenetic trees generated from these amplicons had comparable resolving power to the PsEG30F and PsEG790R pair, albeit with fewer nodes overall. As a consequence, we chose the PsEG30F and PsEG790R primer pair for *Pseudomonas* profiling. The universal 16S rRNA V3V4 primers amplified 100% of the whole genomes and 87.95% of the type strains (owing to 16S genes often being the breakpoint in contigs). As noted previously (20), many of the amplicons are identical across *Pseudomonas* species and hence cannot be used for species resolution (Figure S2).

**Table 1.**
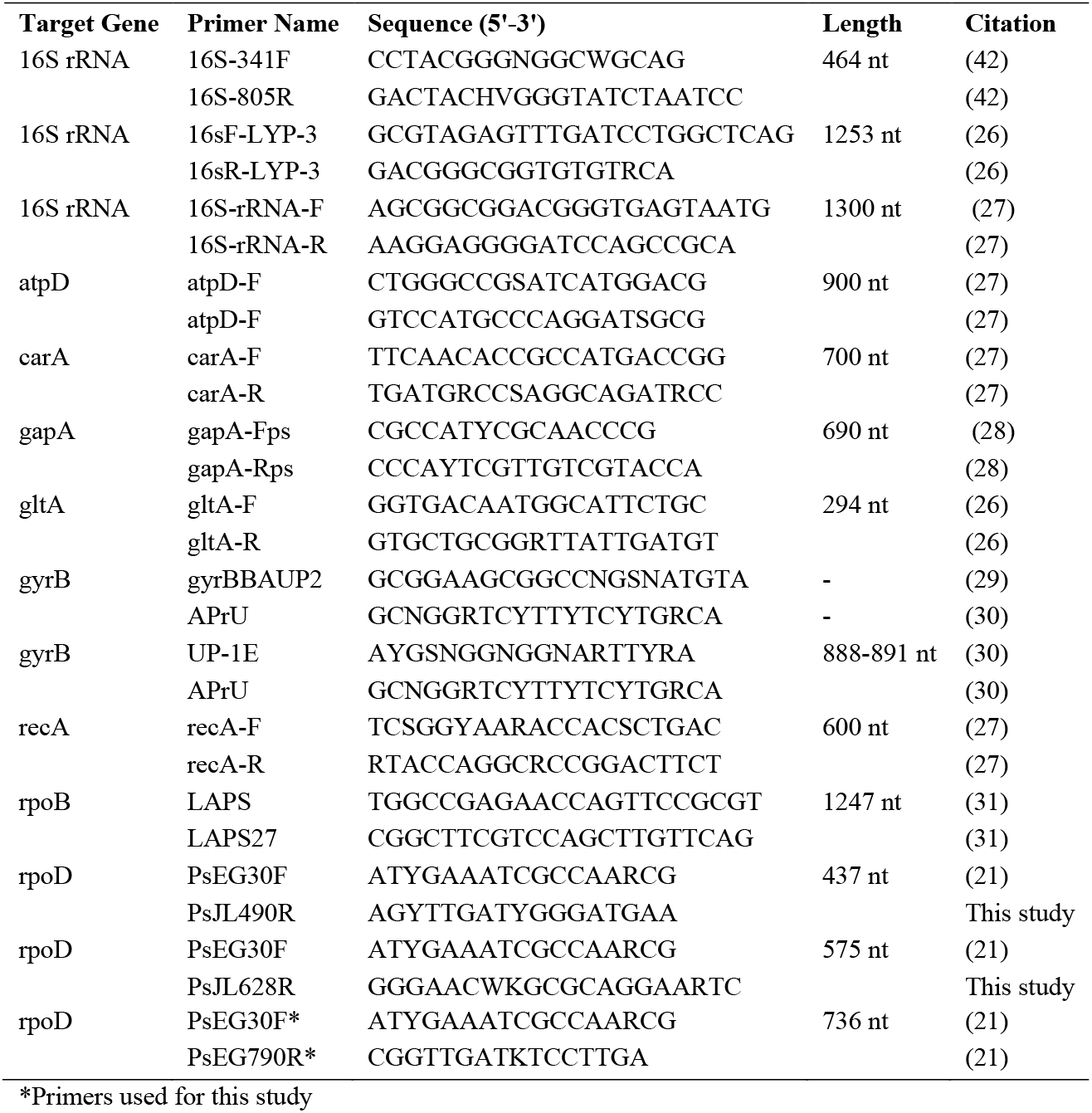
Overview of genes and primers selected for as primary targets for *in_silico_PCR* of 465 *Pseudomonas* strains.

### Synthetic community primer testing

To benchmark the amplicon protocol, a mixture of DNA from sixteen *Pseudomonas* strains was used to test the performance of the candidate *rpoD* primers in comparison with the standard V3V4 16S rRNA gene amplicon sequencing approach. The sixteen strains were selected to challenge the method across the genus by including five groups of *Pseudomonas* species (*P. aeruginosa, P. fluorescens, P. putida, P. stutzeri* and *P. syringae*) and on fine resolution by selecting closely related species especially within subgroups (*P. fluorescens, P. mandelii, P. jessenii*). In contrast to the V3V4-amplicons, PE300 Illumina reads of the *rpoD* amplicons do not overlap and hence cannot be analysed with standard OTU-based methods. To overcome this challenge, each read pair was instead aligned to a custom database of *rpoD* genes using bowtie2 (23).

The *rpoD* amplicon method was able to identify all sixteen strains with abundances close to expected values (Figure 1) and with a very low level of variation across the five technical replicates (Table 2). Of note, two species were underestimated, *P. putida* somewhat (~4% of expected value) and *P. libanensis* severely so (~1% of expected value). In contrast, the V3V4 method erroneously classified the sample composition as being mainly *P. fluorescens* or *P. extremorientalis*, where the latter was not a part of the mixture. Also, low amounts of *P. stutzeri* and *P. aeruginosa* were detected by the V3V4 approach. The beta dispersions - a measure of multivariate variation within groups - of the V3V4 replicates, were 4 times higher than the *rpoD* replicates, although this difference was not significant. The negative controls for both primer sets had low number of reads, later revealed to be common contaminants and adaptors.

**Figure 1.**
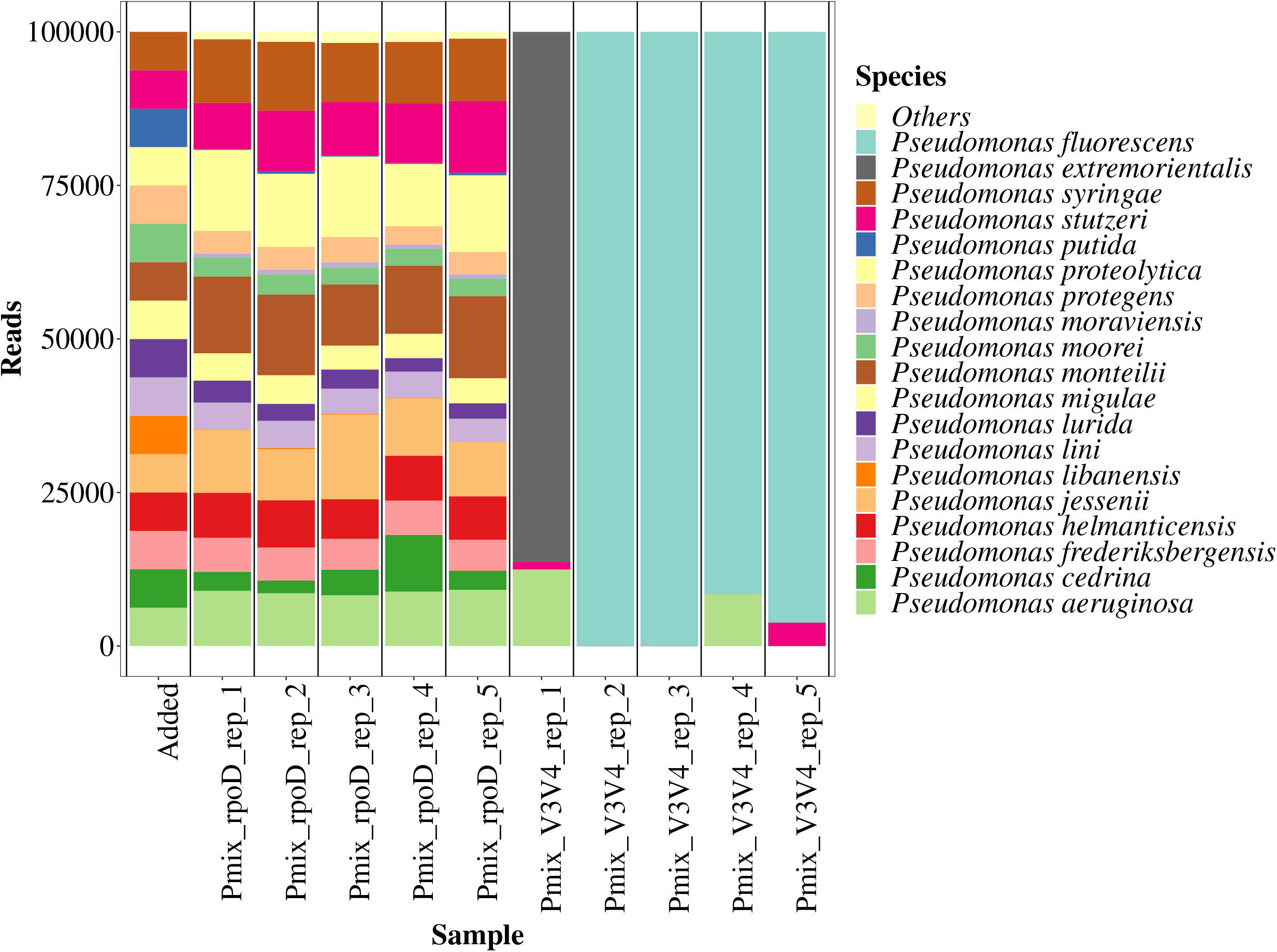
Composition of a defined mixture of DNA from 16 *Pseudomonas* species as analysed by *rpoD* gene amplicon sequencing and V3V4 16S rRNA gene sequencing in comparison to the theoretical composition. Leftmost bar is the assumed theoretical abundances in the defined *Pseudomonas* DNA mixture. Each sample has been normalized to 100.000 reads.

**Table 2.**
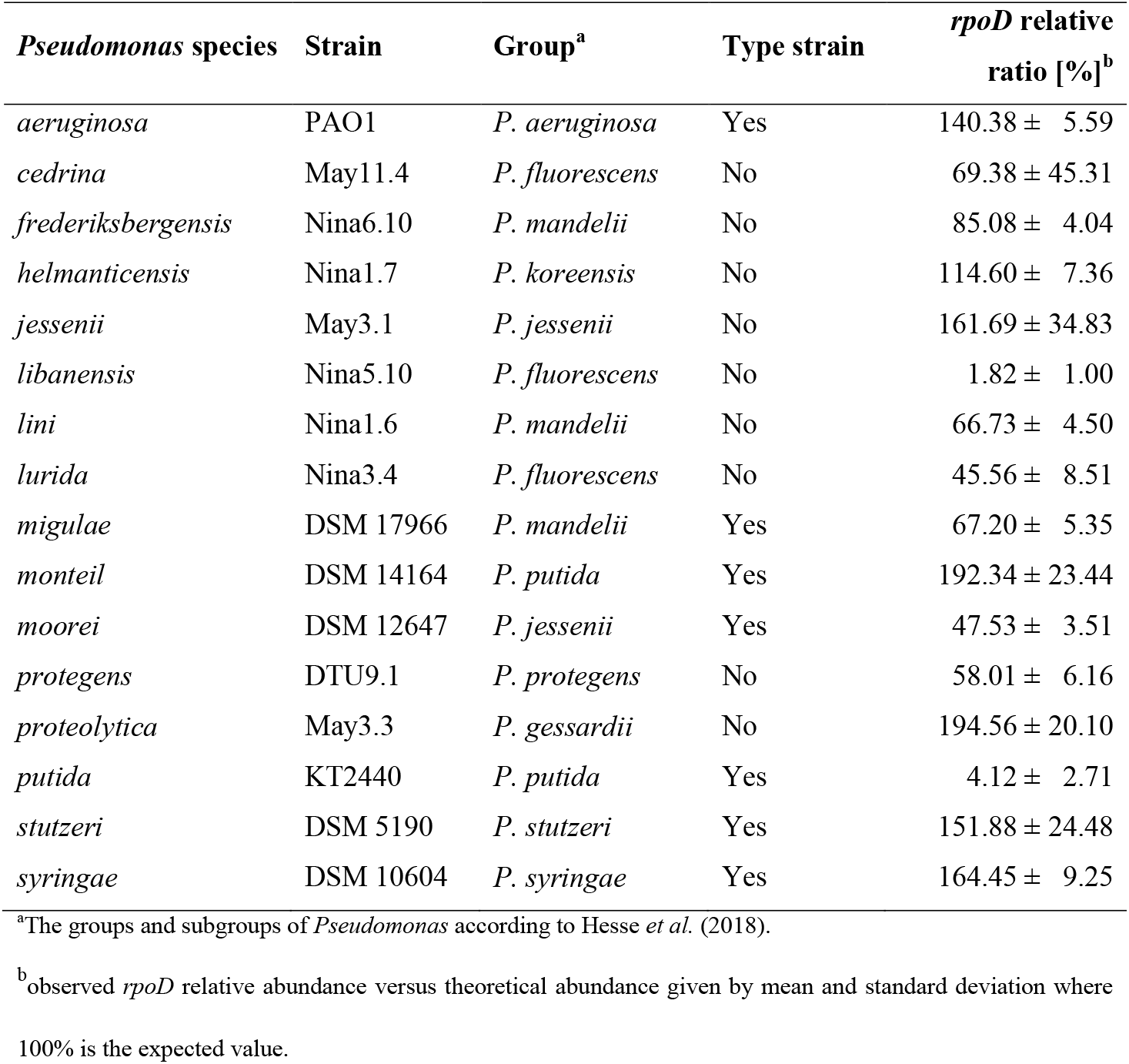
List of *Pseudomonas* species used in the artificial DNA mixture for positive control.

### The microbial and *Pseudomonas* species composition in soil

Soil was sampled from different sites, ranging from grassland to agricultural field soil. A total of 13 soil samples with 5 replicates each (65 samples in total) were analysed and after demultiplexing according to barcodes and primers, a total of 10,446,888 reads were available. Following this, 100,814 *rpoD* read-pairs (201,628 reads) were annotated at the species level for the genus of *Pseudomonas* with sufficiently high confidence (minimum bit score of 10). According to the general purpose metagenomics classification tool Centrifuge (24), the non-*Pseudomonas* reads were approximately 50% PCR/adaptor-artefacts (‘Synthetic constructs’) along with the commonly found contaminant *Bradyrhizobium* also found in the negative control. The overall mean and median values observed for the annotated *rpoD* reads across all samples were 1,558 and 598, respectively, The average number of *Pseudomonas rpoD* reads per sampling site varied between 133.4 (P8) and 4,022.2 (S7). Rarefaction curves revealed uneven saturation in some samples with low read depth (Figure S3), Moreover, we observed that quite few reads were concordantly mapped, this being 5.52%, likely owing to the high level of PCR-artefact non-pseudomonal reads. In addition, less than 0.01% were discordantly mapped.

Using the relative abundances (Figure 2), the performance of the method on the natural samples was investigated across biological replicates. The most abundant species represented the four groups of *P. syringae, P. lutea* (*P. graminis*), *P. putida* (*P. coleopterorum*) and *P. fluorescens* (marked by stars in Figure 2). Within the group of *P. fluorescens*, the five subgroups *P. jessenii* (*P. jessenii, P. moorei, P. umsongensis*), *P. gesardii (P. gesardii, P. proteolytica), P. koreensis (P. baetica, P. helmanticentis, P. moraviensis), P. mandelii (P. frederiksbergensis, P. lini, P. migulae*), and *P. corrugata (P. brassicacearum, P. kilonensis*) were identified. Overall, similar abundances were found in replicate samples. This was confirmed by the non-Metric Multidimensional Scaling (nMDS) analysis (Figure 3), in which biological replicates clustered together although with different degrees of variation. The beta dispersions of the sites were negatively correlated (r=-0.455) to estimated *Pseudomonas* load through qPCR, suggesting that the variation increased as *Pseudomonas* abundance decreased. The two sample sites P8 and P9 had the highest internal variation, likely owing to the low read count in these samples.

**Figure 2.**
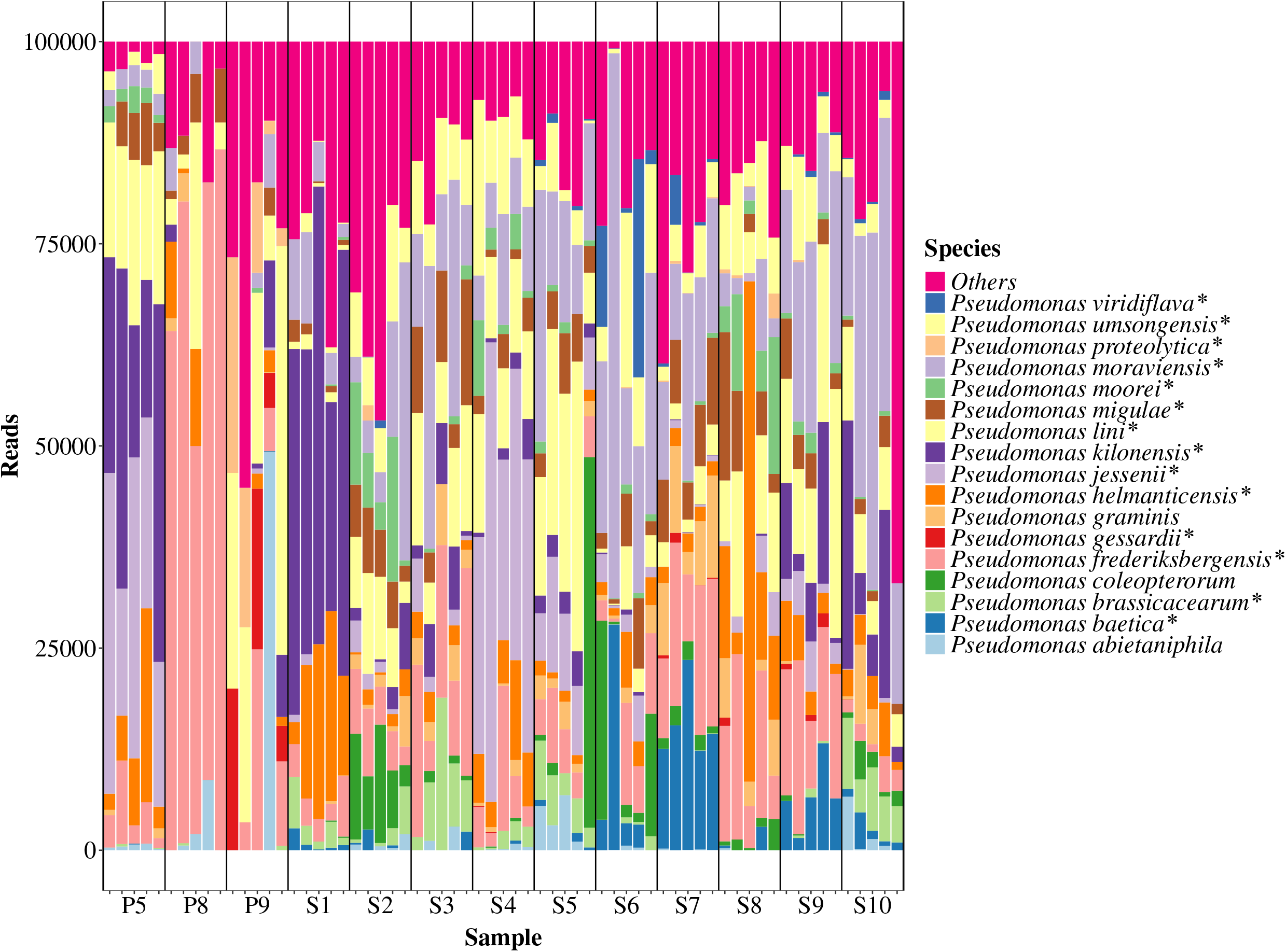
Relative abundances of the 20 most abundant *Pseudomonas* species in 13 soil samples as analysed by *rpoD* amplicon sequencing. Each sample has been normalized to 100.000 reads. Fluorescent species are marked by *. S1: corn, S2: fallow (grass), S3, S7 and S9: wheat, S4: rye, S5: barley, S6: rapeseed, S8: grass seed S10: Lucerne, P5 and P9: pristine short grass, P8: pristine long grass.

**Figure 3.**
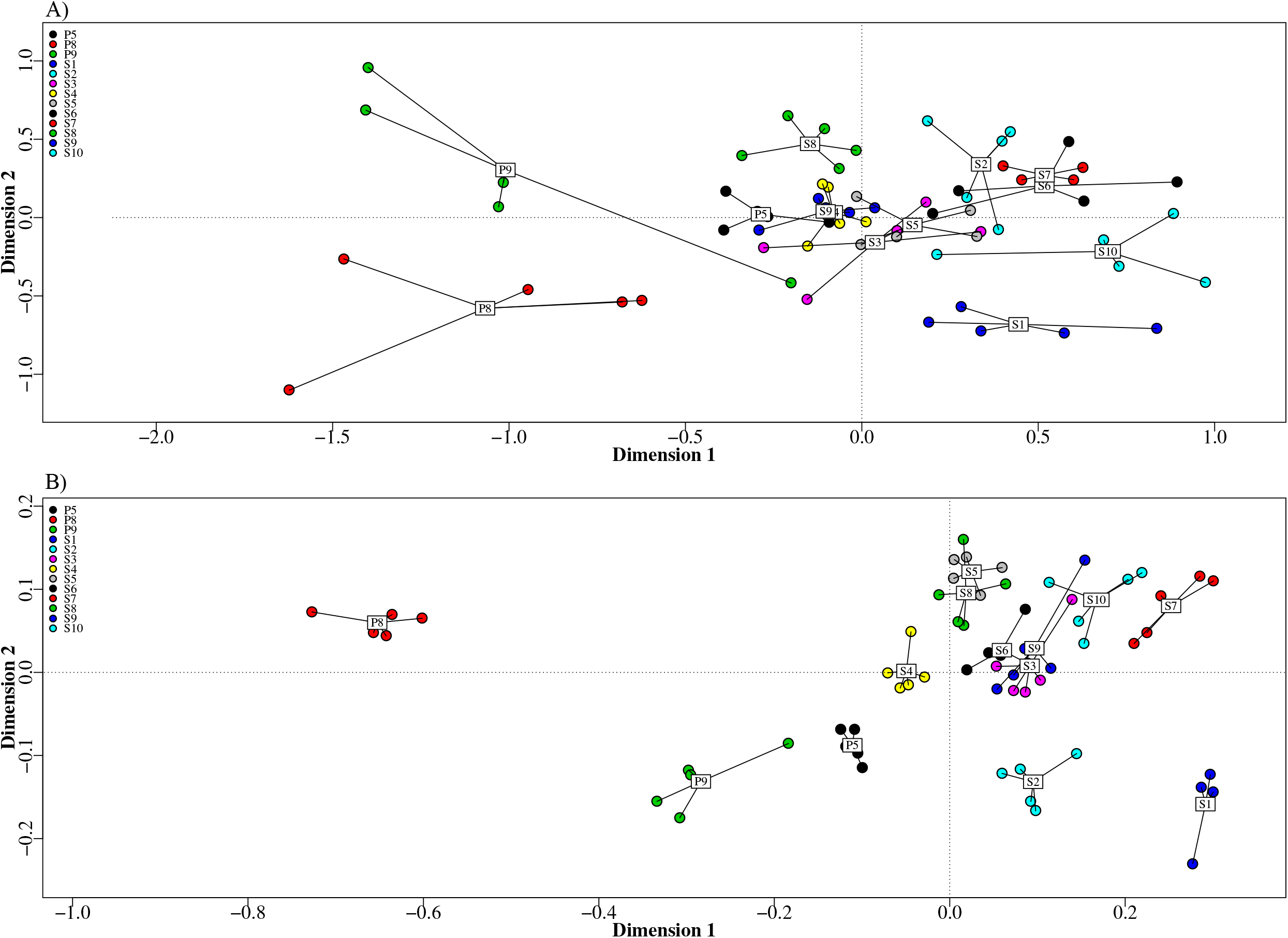
Multivariate analysis by nMDS using Bray-Curtis distances of 13 soil sample sites using amplicon sequencing of A) the *rpoD* gene (stress 0.1883) and B) V3V4-region of the rRNA gene (stress 0.1039). S1: corn, S2: fallow (grass), S3, S7 and S9: wheat, S4: rye, S5: barley, S6: rapeseed, S8: grass seed S10: Lucerne, P5 and P9: pristine short grass, P8: pristine long grass.

The individual sites in the *rpoD* amplicon analysis differed in species composition despite being sampled from similar ecological environments (Figure 2). For instance, the wheat soil samples S3, S7, and S9 had different *Pseudomonas* composition. *P. moraviensis, P. lini, P. helmanticensis* and *P. frederiksbergensis* were common in most or all sampled soils, however, their relative abundances varied between sites.

The standard V3V4 amplicon methodology resulted in a total of 6,947,525 mapped reads with an average of 106,885 per sample. A total of 378 families were identified with the dominating being *Xanthomonadaceae, Sphingomonadaceae, Planctomycetaceae, Chitinophagaceae*, and *Acidobacteria* (group 6) (Figure 4). At the family level, only smaller variations within sample sites were observed, as visually evident in the nMDS (Figure 3), such as the unique presence of *Acidobacteria* (group 1) in the P8 and P9 sites. The average relative abundances of *Pseudomonas* varied from 0.008% (P8) to 0.73% (S8) in the different communities (Table 3). At species level V3V4, typically only identified one or two species in higher relative abundances, and similar to the synthetic communities, these species were assigned to be *P. fluorescens* and *P. aeruginosa*. Each site had, on average, a 70% (P = 0.00004) larger beta-dispersion when profiled for *Pseudomonas* with the V3V4 method compared to the *rpoD* method, and all had at least one sample with a species not found in the other replicates.

**Figure 4.**
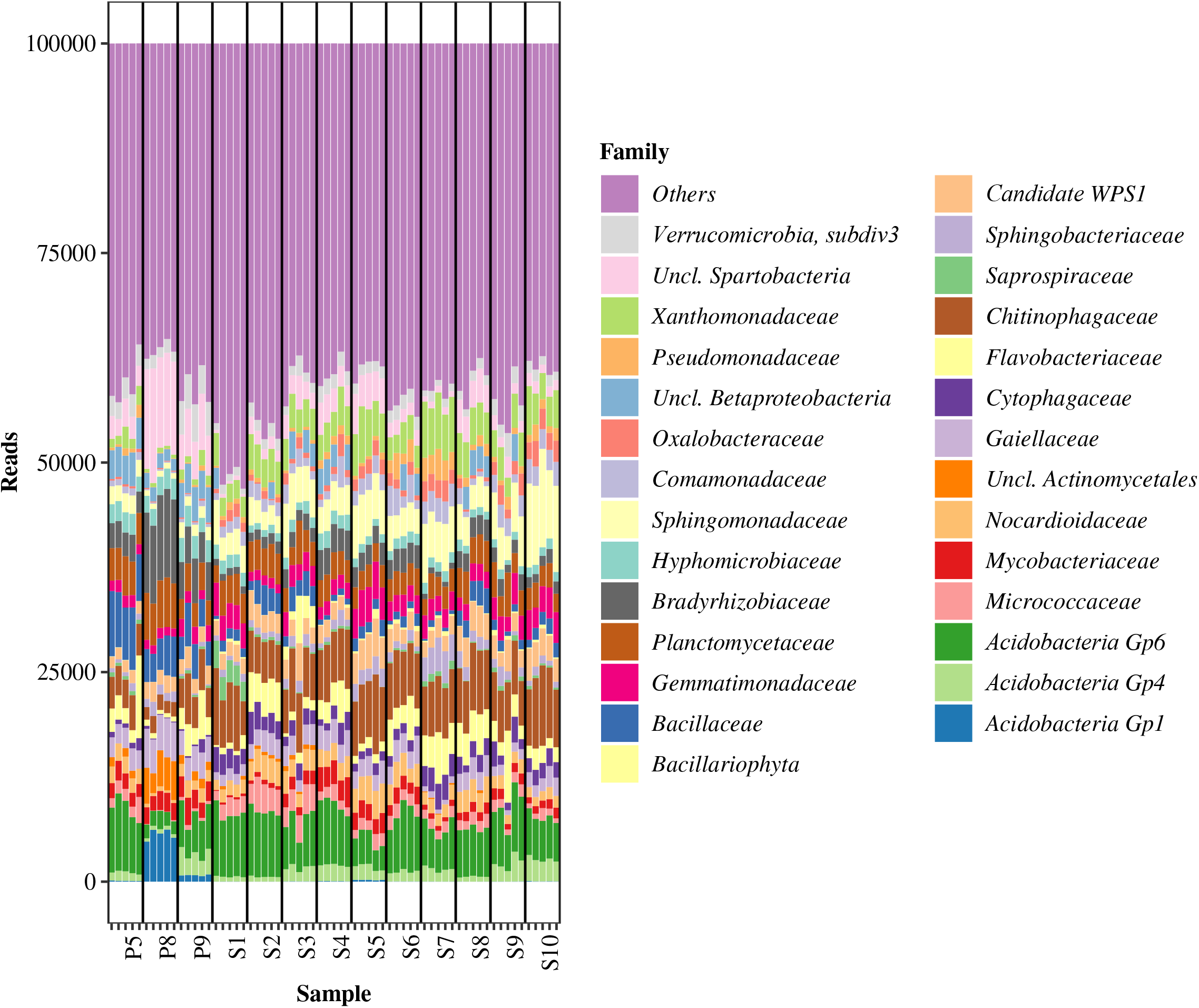
Relative abundances of the 20 most abundant bacterial families in 13 soil samples as analysed by amplicon sequencing of the V3V4-region of the rRNA gene. Each sample has been normalized to 100.000 reads. S1: corn, S2: fallow (grass), S3, S7 and S9: wheat, S4: rye, S5: barley, S6: rapeseed, S8: grass seed S10: Lucerne, P5 and P9: pristine short grass, P8: pristine long grass.

**Table 3.** The average total CFU per site estimated by qPCR and the estimated relative and absolute abundance of *Pseudomonas* species based on V3V4 amplicon sequencing.

| Site | Total bacteria,<br>CFU/g soil | Relative level of<br>*Pseudomonas*, % | Total *Pseudomonas*,<br>CFU/g soil |
| --- | --- | --- | --- |
| P5 | $1.10 \cdot 10^8$ | 0.381 | $4.19 \cdot 10^5$ |
| P8 | $8.13 \cdot 10^7$ | 0.008 | $6.50 \cdot 10^3$ |
| P9 | $5.30 \cdot 10^7$ | 0.027 | $1.43 \cdot 10^4$ |
| S1 | $1.10 \cdot 10^8$ | 0.288 | $3.17 \cdot 10^5$ |
| S2 | $3.37 \cdot 10^7$ | 0.053 | $1.79 \cdot 10^4$ |
| S3 | $3.56 \cdot 10^7$ | 0.021 | $7.48 \cdot 10^3$ |
| S4 | $4.81 \cdot 10^7$ | 0.290 | $1.39 \cdot 10^5$ |
| S5 | $2.21 \cdot 10^7$ | 0.065 | $1.44 \cdot 10^4$ |
| S6 | $2.78 \cdot 10^7$ | 0.649 | $1.80 \cdot 10^5$ |
| S7 | $2.13 \cdot 10^7$ | 0.209 | $4.45 \cdot 10^4$ |
| S8 | $4.29 \cdot 10^7$ | 0.731 | $3.14 \cdot 10^5$ |
| S9 | $4.27 \cdot 10^7$ | 0.126 | $5.38 \cdot 10^4$ |
| S10 | $2.73 \cdot 10^7$ | 0.276 | $7.53 \cdot 10^4$ |

A standard curve comparing Ct-values from qPCR and cell numbers was generated using pure cultures of *P. moorei* and *Bacillus subtilis* and used to estimate total rather than relative abundance. The average total CFU/g soil per sample site (Table 3) ranged from 2.1·10^7^ (S7) to 1.1·10^8^ (S1 and P5). The average number of *Pseudomonas* CFU/ g soil was calculated by multiplication with the relative abundance of *Pseudomonas* found in the V3V4 amplicons (Table 3) and ranged from 6.5·10^3^ (P8) to 4.2·10^5^ (P5) CFU/ g soil.

### Isolation and identification of presumptive *Pseudomonas*

The *rpoD* amplicon culture-independent method for *Pseudomonas* species profiling was compared to cultivation-based profiling. Colonies were isolated from each of the sites on King’s Agar B^+++^ commonly used for *Pseudomonas* isolation (25), 10 from each of the S1-S10 sites and 30 for the grassland samples P5, P8 and P9. The isolates were taxonomically classified by Sanger sequencing of part of the *rpoD* gene amplified with primers PsEG30F and PsEG790R and alignment to the *rpoD* database used for the amplicon analysis. In the grassland samples, 88 of the 90 isolates were classified at a species level. Of particular interest were the species belonging to the *P. fluorescens* group to which encompassed 94% (83/88) of the isolates. This agrees with culture-independent profiling of P5 and P8 in which 97% (P5) and 93% (P8) of the *rpoD* reads were assigned to species in the *P. fluorescens* group. In P9, the fraction of species in the *P. fluorescens* group was lower (52%) which was predominantly due to a high fraction of *P. abietaniphila* species (39.5%) which was not observed by cultivation (Figure 5 & Figure S4).

**Figure 5.**
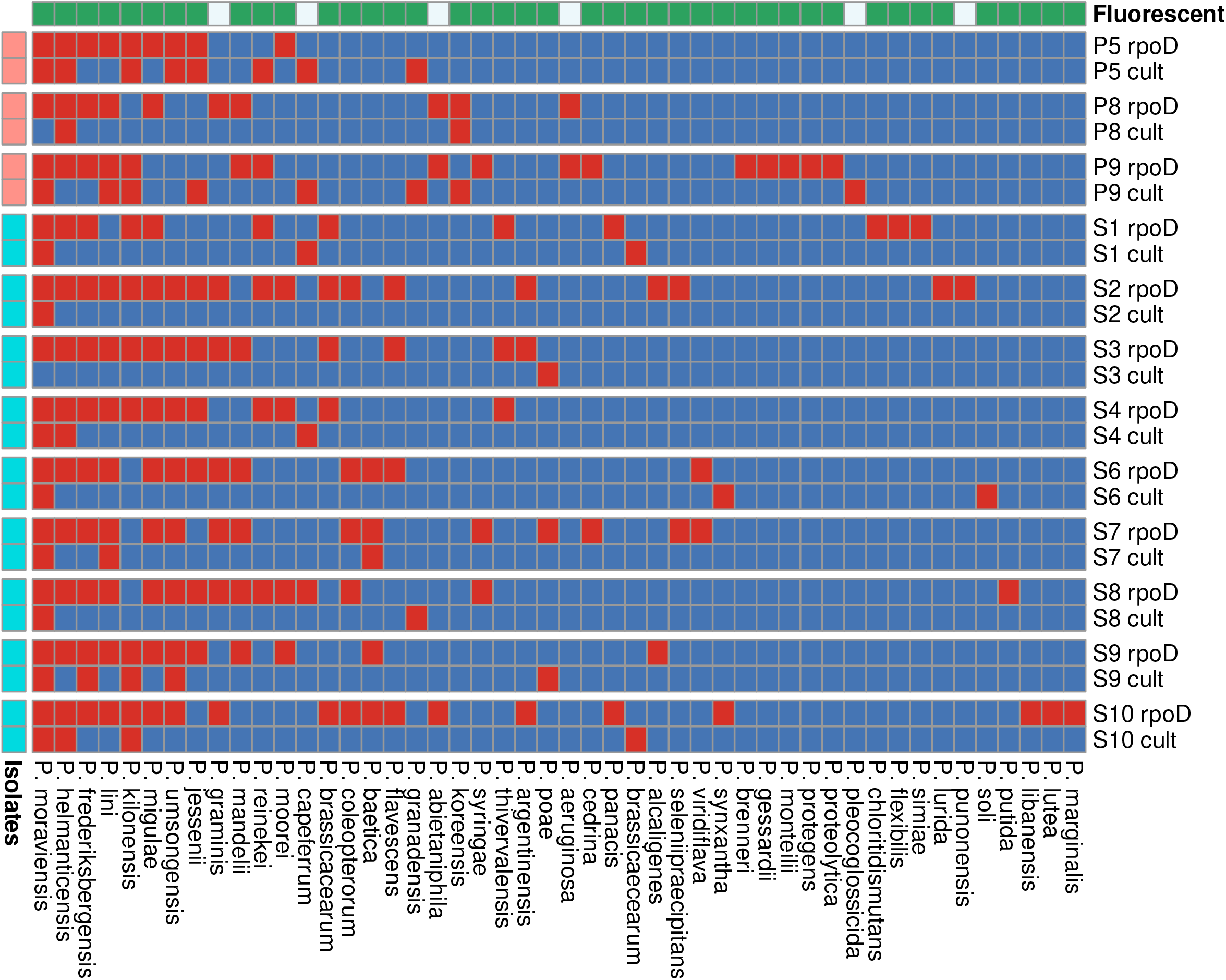
Heatmap of presence (red) and absence (blue) of *Pseudomonas* species identified across sites by the *rpoD* amplicon method and the cultivation approach for each of the sampled sites. 30 isolates were sampled from each of the sites P5, P8 and P9, and 10 from the sites S1-S10 (left-most colour annotation). Fluorescent species are highlighted by green in the upper colour annotation. Row labels denoted with –rpoD and –culture denote analysis by rpoD sequencing and culture, respectively.

The *rpoD* amplicon method identified more unique species than the cultivation method (Figure 5) and species belonging to particular groups (i.e. the *P. lutea* group) and sub-groups (i.e. the *P. mandelii* and *P. gessardii* subgroups of *P. fluorescens*) were almost exclusively identified by the amplicon method but not with the cultivation-based method. The same pattern emerged in the lesser studied sites, S1-S10, where *rpoD* profiling identified between 12 and 19 species compared to the cultivation approach which found between zero and five.

## DISCUSSION

Global food demand is growing and since petrochemical-based industrial farming is unlikely to be sustainable for future generations, there is an urgent need for novel and sustainable biocontrol agents. Species of the *Pseudomonas* genus are promising as plant biocontrol agents and since beneficial traits are typically linked to particular species, we developed a high-throughput method for metataxonomic assignment of these species in natural microbiomes. The method correctly identified all species of a *Pseudomonas* mock community. In soil samples, the *rpoD* amplicon sequencing allowed a much higher degree of *Pseudomonas* species differentiation than both traditional 16S rRNA V3V4 sequencing and culturing. The *rpoD* profiling enables quick identification and prioritisation of soils with specific *Pseudomonas* communities for further analysis and culturing.

A total of fourteen primer pairs targeting the genus *Pseudomonas* (26–31) were examined using *in silico* PCR. The primer pair PsEG30F and PsEG790R (21) outperformed all other pairs and in further analysis of the *rpoD* genes from 465 *Pseudomonas* species, two alternative forward primers were identified. However, the PsEG30F and PsEG790R had superior performance and was ultimately selected for further testing. Multiple studies (20, 21, 26, 32, 33) have shown that the *rpoD* gene is a good candidate for identification at the species- and strain level for the genus of *Pseudomonas*, especially compared to the 16S rRNA gene (20). The second-best candidate was identified as the *gap* primers of Sarkar and Guttman (28). However, these primers also amplified *non-Pseudomonas in vitro* (data not shown) and were therefore discarded. The PsEG30F and PsEG790R amplicons were adapted to an Illumina system, which unfortunately has too short of a read length to overlap, necessitating a new bioinformatic pipeline drawing inspirations from annotation of RNA-seq data, as well as a database based on the genomes from Hesse *et al*. (2018) (15).

When testing the method on a known mixture of pseudomonads, *P. libanensis* and *P. putida* were underestimated. Through *in silico* investigation *P. libanensis* was poorly targeted by the primers implying poor amplification efficiency (data not shown). The *rpoD* genes of *P. putida* KT2440 (in the mixture), *P. putida* NBRC14164 (in database), and *P. monteilii* DSM14164 (in database) were compared in a phylogenetic tree (Figure S5), and although the three species are extremely closely related, KT2440 and DSM14164 are nearly identical and likely to overlap in the investigation. In the future, the *rpoD* database could be expanded to contain more strains for each species to give a wider coverage for such fringe cases. Such an addition could also lead to a strain level differentiation in future studies. Alternatively, this occurrence could also indicate that *P. putida* and *P. monteilii* generally are very closely related and difficult to separate. According to Hesse et *al*. (15), the two species are also extremely closely related based on protein phylogeny of 100 gene orthologues.

The V3V4 amplicon data for the *in vitro* DNA sample predominantly identified one species in the sample, *P. extremorientalis* or *P. fluorescens*, both of which belong to the subgroup *P. fluorescens* (15). This was also seen in Mulet *et al*. (20), where species of the *Pseudomonas* genus at most had 2% dissimilarity in the 16S rRNA gene, this gene did not allow a species level resolution.

The *rpoD* amplicon methodology was used to profile the *Pseudomonas* population in soil samples. The relative abundances across replicates were very similar, yet, some variation was observed, which could be caused by spatial differences within the soil sampled (34). Replicate variance was associated with low *Pseudomonas* load as a negative correlation (r= - 0.455) was observed between the multivariate variance of the replicates (e.g. the beta-dispersion) and the observed *Pseudomonas* CFU/g soil. To our knowledge the closest non-16S rRNA gene based amplicon study of *Pseudomonas* is Sánchez *et al*. (22), where an *rpoD* amplicon methodology was used to identify *Pseudomonas* species in a water sample.

However, a direct comparison is difficult, since Sánchez *et al*. (22) analysed a single river sample with no replicates and used 454 sequencing and a blastn similarity search followed by OTU clustering rather than Illumina sequencing and bowtie-mapping as in our study. The use of the 454 system, discontinued in 2016, results in longer (300-600bp) single end reads, thus avoiding the issues with non-overlapping reads, although they are also too short to cover the entire amplicon. Use of single-end reads allows for analytically simpler OTU-based pipelines, but discards the phylogenetically import paired end information of our approach, and hence has lower sensitivity. Sánchez *et al*. (22) assigned 10.8% (716 sequences) of the *rpoD* gene sequences to one of twenty-six species in their database. Using the genomes of Hesse *et al*. (15), the database now includes 166 *Pseudomonas* species, three of which are subspecies. Many of the *rpoD* reads were not mapped to our *Pseudomonas* database, which is likely due to the high stringency in our alignment approach and artefacts stemming from the low template PCR reaction, e.g. when *Pseudomonas* is in low abundance compared to other bacteria. Optimisation of the PCR-protocol may alleviate this.

The microbial composition as determined by V3V4 16S rRNA amplicon sequencing was highly similar across biological replicates, with small differences between the sample sites. The three wheat-associated sites (S3, S7 and S9) clustered, but had less overlap than a cluster with sites S3, S6 and S9 which all had different vegetation. Microbial communities in agricultural soils are influenced by physiochemical properties of the soil, the growth condition of the crops, the individual plant genotype, and/or the evolution of the microbial communities over a multitude of seasons and the present study did not have access to such metadata that potentially could explain differences. The major outlier of the sites was the P8 site, mainly due to the high content of *Acidobacteria* (group 1), suggested to correlate with Cu and Mn concentrations (35), coinciding with a high relative abundance of *P. frederiksbergensis*.

The resolution at the species level was compared between the *rpoD* and the V3V4 amplicon sequencing methods. The *Pseudomonas* species resolution based on the latter was lower than found by the *rpoD* amplicon sequencing, which was consistent with the control experiment using 16 known species of pseudomonads. This is most likely a combination of the low dissimilarity in the 16S rRNA gene, annotation method and the database. It is important to note the usefulness of V3V4 rRNA gene amplicon sequencing as a tool to determine the overall composition of the prokaryotic community. Different microbial communities were observed across the different soils, some of which even have the same plant host. The *rpoD* amplicon methodology did not achieve a community level resolution and is therefore best used in combination with standard 16S profiling to achieve full profiling of soil.

The *rpoD* profiling was compared to cultivation of *Pseudomonas* species from three soil samples and provided the same groups or sub-groups, however, the *rpoD* amplicon sequencing method was able to identify more unique species than was found by cultivation. In particular, species belonging to the *P. lutea* group and the *P. mandelii* and *P. gessardii* subgroups of *P. fluorescens* were captured by the *rpoD* amplicon method but not by cultivation. The nutrient content of the isolation medium can influence the recovery of the *Pseudomonas* diversity from environmental samples (19). King’s B agar medium (36) is a nutrient rich medium for *Pseudomonas* isolation, and while this is an effective and commonly used cultivation method, it is possible that a larger number of *Pseudomonas* species could have been cultivated using a wider range of cultivation media. Overall, the *rpoD* amplicon methods can be used to find soil rich in *Pseudomonas* species and identify samples rich in potential beneficial or useful pseudomonads.

A few species were exclusively found by the culture-independent approach, and some of these are promising as bio-inoculants for plant protection (Figure 5). For example, strains of *Pseudomonas frederiksbergensis* (from the *P. mandelii* sub-group), which were found in all *rpoD*-profiles, but only once by cultivation (in sample S9), are effective bio-inoculants for enhancing cold stress tolerance in plants (37). In addition, *Pseudomonas abietaniphila* (from the *P. lutea* group), which was found in three *rpoD* profiles (P8, P9 and S10), but not in any cultivations, can suppress plant diseases caused by *Botrytis cinerea* by degradation of oxalate produced by the fungi (38). Also, we have recently shown that the genes for biosynthesis of the anti-fungal compound thioquinolobactin are rarely found, but tightly linked to biocontrol strains within the *P. gessardii* subgroup of *P. fluorescens* (39). Here, we find such species in P9 only with the cultivation-independent method.

With the *rpoD* amplicon approach, it is possible to profile and prioritise samples for intense cultivation of strains that produce specific bioactive metabolites for biocontrol or exhibit other plant beneficial traits.

## CONCLUSION

In this study, an *rpoD* gene amplicon-based technique to differentiate species within the genus *Pseudomonas* was developed. The method can differentiate individual species far beyond what traditional 16S rRNA gene amplicon sequencing can and is proposed as a new standard for high throughput profiling of *Pseudomonas* in environmental microbial communities.

## MATERIAL AND METHODS

### *In silico* investigations of *Pseudomonas* species

A *Pseudomonas* genome collection consisting of the 165 genomes from Hesse et al. (15) as well as 465 complete genome sequences of *Pseudomonas* was downloaded from NCBI (downloaded 18-02-2019). Using an *in silico* PCR algorithm (*in_silico_*PCR, https://github.com/egonozer/in_silico_pcr) (one mismatch, one deletion/insertion), fourteen primer sets targeting nine different genes from previous studies (Table 1) were evaluated based on 1) the proportion of *Pseudomonas* genomes amplified, 2) how well the amplicons followed their given phylogenetic classification and 3) the proportion of *non-Pseudomonas* genomes amplified (supplementary Table S1). The *in silico* PCR products were aligned with MUSCLE v3.8.1551 (40) and a phylogenetic tree was generated with FastTree v2.1.10 (41) from the alignment. For phylogenetic evaluation, the output tree was qualitatively compared to whole-genome based tree of Hesse *et al*. (15). A custom database was built by using the PsEG30F / PsEG790R primers on the *rpoD* genes of the type strains included in Hesse *et al*. (15). An issue encountered in the *in silico* analysis was the poor annotations of uploaded genomes where we found multiple instances of incorrectly annotated genomes, which we corrected by selecting the *rpoD-gene* of outliers and re-identify them according to Hesse *et al*. (15) type strain genomes.

### *Pseudomonas* strains

Sixteen different *Pseudomonas* species were used in the *in vitro* testing of the *rpoD* amplicon (Table 2). Seven were Type Culture collection strains and nine were isolates obtained from on-going projects in our laboratory. They were identified to species level by Sanger sequencing of the *rpoD* gene as described above. The strains were grown in 10 mL Luria-Bertani (LB; Lennox; Carl Roth GmbH + Co. KG, Karlsruhe, Germany) broth overnight at 30°C with aeration (shaking, 200 rpm).

### Soil samples

Bulk soil samples were collected from thirteen sites in mid-August 2019 close to harvest season. The samples were collected by scooping root-associated soil into a sterile Falcon tube. The sites were distributed across Zealand, Denmark, and included different types of vegetation and produce; ten samples of field soil were collected including corn (S1), fallow (grass; S2), wheat (S3, S7 and S9), rye (S4), barley (S5), rapeseed (S6), grass seed (S8) and lucerne (S10). In addition, three samples of pristine grass land were collected including short (P5 and P9) and tall grass (P8). The soil samples were stored at 5°C for a maximum of two weeks prior to analyses.

### Isolation of *Pseudomonas* from soil samples

Soil was sieved (4.75 mm X 4.75 mm grid) and mixed with 0.9% NaCl in a 1 (g):9 (ml) ratio and 10-fold serial diluted. Dilutions were plated on 1/4-diluted King’s B^+++^ Agar plates (Sigma-Aldrich) supplemented with 13 mg/L chloramphenicol, 40 mg/L ampicillin, 100 mg/L cycloheximide (25)).. The plates were incubated at 30°C for 5 days. The plates were examined under UV-light after 2 and 5 days to reveal fluorescent colonies. Up to thirty colonies from the P5, P8 and P9 sites as well as 10 from the S1-S10 sites were streaked on LB Agar plates and incubated at 30°C for 2 days. The colonies were selected based on fluorescence and distinct colony morphology.

### DNA extraction from pure cultured *Pseudomonas* and soil samples

For identification of *Pseudomonas* isolated from soil, DNA from each bacterial colony was extracted by boiling in demineralised H_2_O (dH_2_O) at 99°C for 15 min. For soil samples and selected *Pseudomonas* strains, Genomic DNA (gDNA) was extracted with a DNAeasy^®^ Powersoil^®^ Kit (Qiagen, Hilden, Germany) according to manufacturer’s instructions. The extractions of gDNA from soil were done in five biological replicates for each site. As a negative control, 250 μL sterile dH_2_O was extracted for gDNA using the same methodology. The gDNA was stored at −20°C. The DNA extraction of soil was done at the latest two days after cultivation of soil *Pseudomonas*.

### Identification of *Pseudomonas* isolates from soil samples

*Pseudomonas* species isolated from soil were identified by full length sequencing of the *rpoD* gene. The 25 μL PCR reaction mixture was; 10.6 μL Sigma Water, 12.5 μL 2x TEMPase, 0.8 μL forward primer (10 μM; PsEG30F; 5’-ATY GAA ATC GCC AAR CG-3’), 0.8 μL reverse primer (10 μM; PsEG790R; 5’-CGG TTG ATK TCC TTG A −3’), and 0.3 μL template DNA. The PCR program was: 1) 1 cycle of 95°C for 15 min, 2) 30 cycles of a) 95°C for 30 s, b) 51°C for 30 s, and c) 72°C for 30 s, and 3) 1 cycle of 72°C for 5 min (21). PCR products were sequenced at Macrogen Europe (Amsterdam, the Netherlands). The *rpoD* sequences were classified using BLASTN against the custom built *rpoD* gene database.

### Defined *in vitro* DNA mixture for *Pseudomonas* profiling

As a positive control, a defined *Pseudomonas* DNA mixture was made as an equimolar mixture of individual extractions of the strains in Table 2 as measured by Nanodrop (Denovix DS-11; Saveen & Werner AB, Linhansvãgen, Sweden). The equimolar mixture was based on DNA concentrations.

### Amplicon preparation, purification and sequencing

Amplicons were prepared by amplifying DNA using barcoded primers (Table S2). The five biological replicates of each soil site, five technical replicates of the *in vitro* DNA mix (positive control) and the negative control all were amplified using both the *rpoD*-specific primers and primers targeting the V3V4 region of the 16S rRNA gene. Each sample used identical barcodes across both primer sets (Table S2) and Illumina adaptors for the two setups. For the amplification of *rpoD* genes, 25 μL PCR reaction was mixed as 10.15 μL Sigma Water, 12.5 μL 2x TEMPase, 0.8 μL forward primer (10 μM; Barcoded PsEG30), 0.8 μL reverse primer (10 μM; Barcoded PsEG790), 0.25 μL MgCl_2_ (25 mM), and 0.5 μL Template DNA were used. The PCR program was as follows 1) 15 min at 95°C, 2) 40 cycles of a) 30 s at 95°C, b) 30 s at 51°C, and c) 30 s at 72°C, and 3) 5 min at 72°C. The amplicons were stored at −20°C until purification.

For the amplification of V3V4 regions, 25 μL PCR reaction was missed as; 10.6 μL Sigma Water, 12.5 μL 2x TEMPase, 0.8 μL forward primer (10 μM; Barcoded 341F; 5’-CCT ACG GGN GGC WGC AG-3’), 0.8 μL reverse primer (10 μM; Barcoded 805R; 5’-GACTACHVGGGTATCTAATCC-3’), and 0.3 μL Template DNA were used (42). The PCR program was as follows 1) 15 min at 95°C, 2) 30 cycles of a) 30 s at 95°C, b) 30 s at 60°C, and c) 30 s at 72°C and 3) 5 min at 72°C. The amplicons were stored at −20°C until purification.

The amplicons were purified using an Agencourt AMPure XP kit (Beckman Coulter, Brea, CA, USA) following the manufacturer’s instructions. The products were eluted in tris(hydroxymethyl)aminomethane (Tris; 10 mM, pH 8.5) buffer. After purification, the PCR products were equimolar pooled together.

The amplicon pools were delivered at the CfB NGS Lab (Novo Nordisk Foundation Center for Biosustainability, DTU, Kongens Lyngby, Denmark) for sequencing on an Illumina MiSeq 300PE platform (MiSeq Reagent Kit v3; PE300).

### Enumeration of soil bacteria

The number of cells in each soil site were quantified using quantitative PCR (qPCR). The qPCR targeted the V3 region of the 16S rRNA gene using the primers 338F and 518R (43). A 20 μL PCR reaction was mixed as follows: 5.2 μL Sigma Water, 10 μL Luna Universal qPCR Master Mix (New England Biolabs Inc., Bionordika Denmark A/S, Denmark), 1.4 μL of each primer (10 μM), and 2 μL Template DNA. The accompanied instruction for the qPCR programme was followed. A standard curve relating cycle thresholds (Ct) to CFU/g soil was prepared by combining CFU/g versus Ct for *Bacillus subtillis* ATCC 6051 and *Pseudomonas moorei* DSM 12647 (R^2^ = 0.86, E = 174.5%). ATCC 6051 and DSM 12647 were incubated O/N in 5 mL LB broth at 30°C with aeration. At OD_600_ approximately 1 (circa 24 hours of growth), DNA was extracted from the cultures and further diluted. The standard curves were prepared in biological duplicates.

### Processing the V3V4 and *rpoD* amplicons

The V3V4-amplicons were cleaned, merged, quality filtered, and chimera-checked before quality-aware clustering at 99% similarity and mapping against the RDP-II SSU (Cole et al., 2005) database (v. 11.5) using the BION-meta software (Danish Genome Institute, Aarhus, Denmark). For the *rpoD*-amplicons (PsEG30F / PsEG790R), the BION-meta software (Danish Genome Institute, Aarhus, Denmark) was used to demultiplex the amplicons. The fastp function (44) was used for quality filtering. Since the paired reads do not overlap, clustering was avoided and instead each read pair was aligned to the custom database of *rpoD* genes using bowtie2 (23). The resulting SAM-file was then filtered for only concordant pairs mapped with a quality of > 10 using samtools (45). Data for both sets of amplicons was normalized to 100,000 reads for each sample before further analysis. Centrifuge was used for profiling of *non-Pseudomonas* reads using the p+h+v database (24).

### Statistics

The amplicon sequencing data for both *rpoD* and V3V4 were analysed with Non-metric multidimensional scaling (nMDS) to compare the diversities between the replicates and sample sites. To determine the multivariable variation within groups, the beta dispersion was calculated using R v3.6.2 package vegan, default settings and tested using a Mann-Whitney U test. Multiple distances were evaluated for robustness and the Bray-Curtis distance was chosen since this distance metric had the best trade-off in terms of separation of sites and stress of the nMDS.

### Data availability

The raw, amplicon sequencing data of the Illumina sequencing is available at the Sequencing Read Archive (SRA) as bioproject PRJNA613913. Code for both *in silico* primer analysis and the bioinformatic classification pipeline is available at https://github.com/mikaells/PseudomonasRPOD.

## ACKNOWLEDGEMENTS

We thank local farmers and governmental agencies, whom provided sites for soil collection, Heiko T. Kiesewalter for assistance with sampling and providing *Bacillus subtillis* ATCC 6051, and the laboratory technicians Jette Melchiorsen, Sophia Rasmussen and Susanne Koefoed for support.

## LEGENDS FOR SUPPLEMENTARY TABLES AND FIGURES

**Figure S1:** Gel electrophoresis of PCR products generated from members of the synthetic community using A) PsEG30F / PsEG790R, B) PsEG30F / PsJL490R, and C) PsEG30F / PsJL628R. Contrasts has been enhanced. Ladders are 50 bp GeneRuler (ThermoFisher Scientific, Vilnius, Lithuania).

**Figure S2**. Phylogenetic tree of the *in sillico* PCR amplicons generated from the 16S rRNA gene using V3V4 primers. The PCR products were clustered at 97% similarity and aligned with MUSCLE v3.8.1551 (40) and a phylogenetic tree was generated with FastTree v2.1.10 (41) from the alignment. Tips are coloured as clades generated by TreeCluster with similar colours denoting highly similar or identical amplicons.

**Figure S3:** Alpha diversity of the *Pseudomonas* population as described using the rpoD-methodology. A) rarefaction curves and B) Chao1 diversity.

**Figure S4.** Number and identity of *Pseudomonas* species isolated by cultivation from the 13 soil sample sites.

**Figure S5**. Phylogenetic tree of the *in sillico* PCR amplicons generated from the *rpoD* gene of *P. putida* KT2440 and four closely related species. The PCR products were aligned with MUSCLE v3.8.1551 (40) and a phylogenetic tree was calculated with FastTree v2.1.10 (41) from the alignment.

**Table S1**: Genomes of non-*Pseudomonas* used to test *Pseudomonas* gene primer specificity profiling.

**Table S2**: List of barcodes tagged on primers for Illumina sequencing of amplicons of the *Pseudomonas* rpoD gene.

**Table S3**. *In silico* PCR amplification of 465 complete *Pseudomonas* genomes and 24 non-*Pseudomonas* genomes using 14 different primer pairs.

